## Supplementary figures and images for "Sex chromosome evolution in beetles"

### Supplementary Figure 1

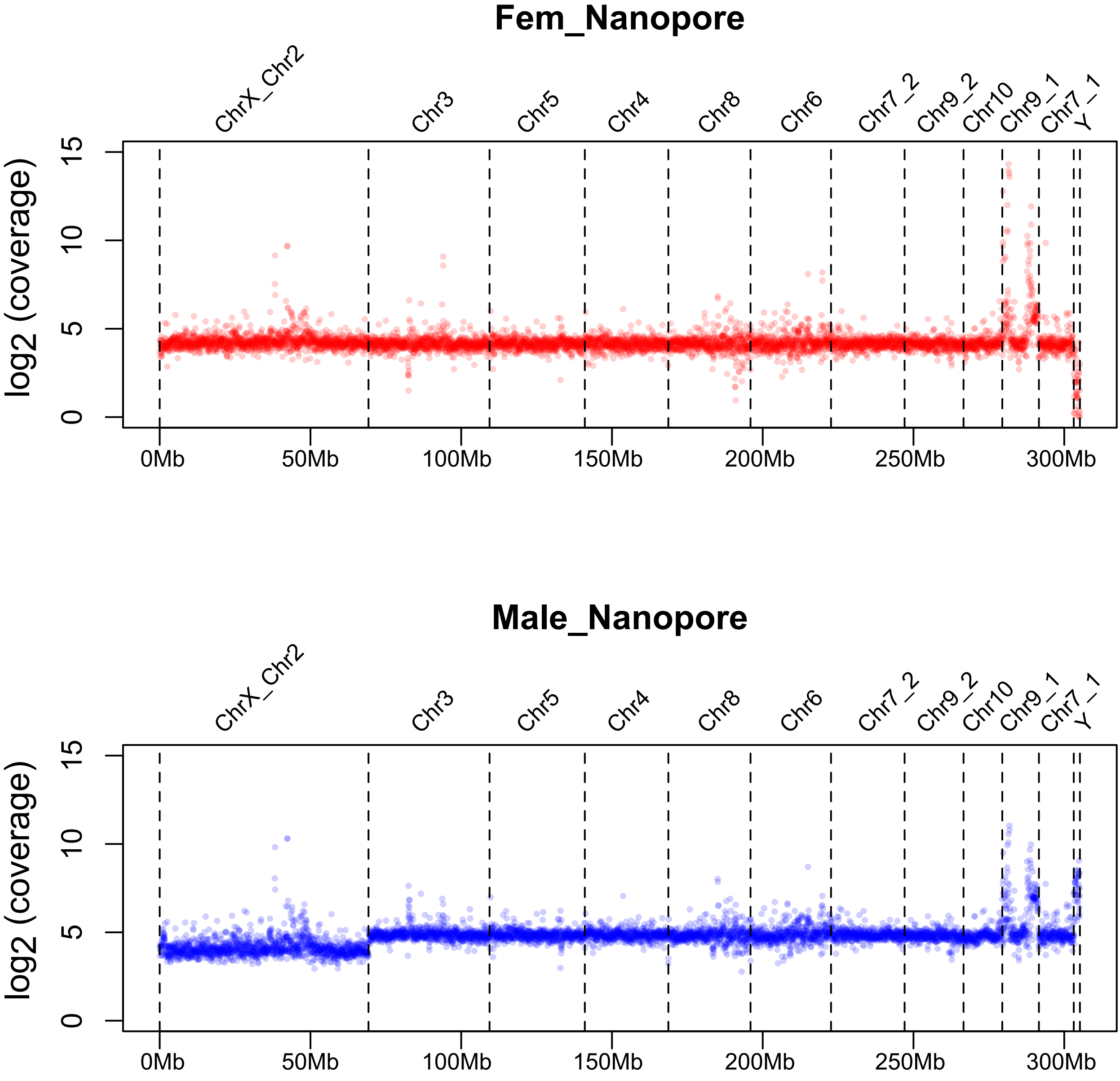

### Supplementary Figure 2

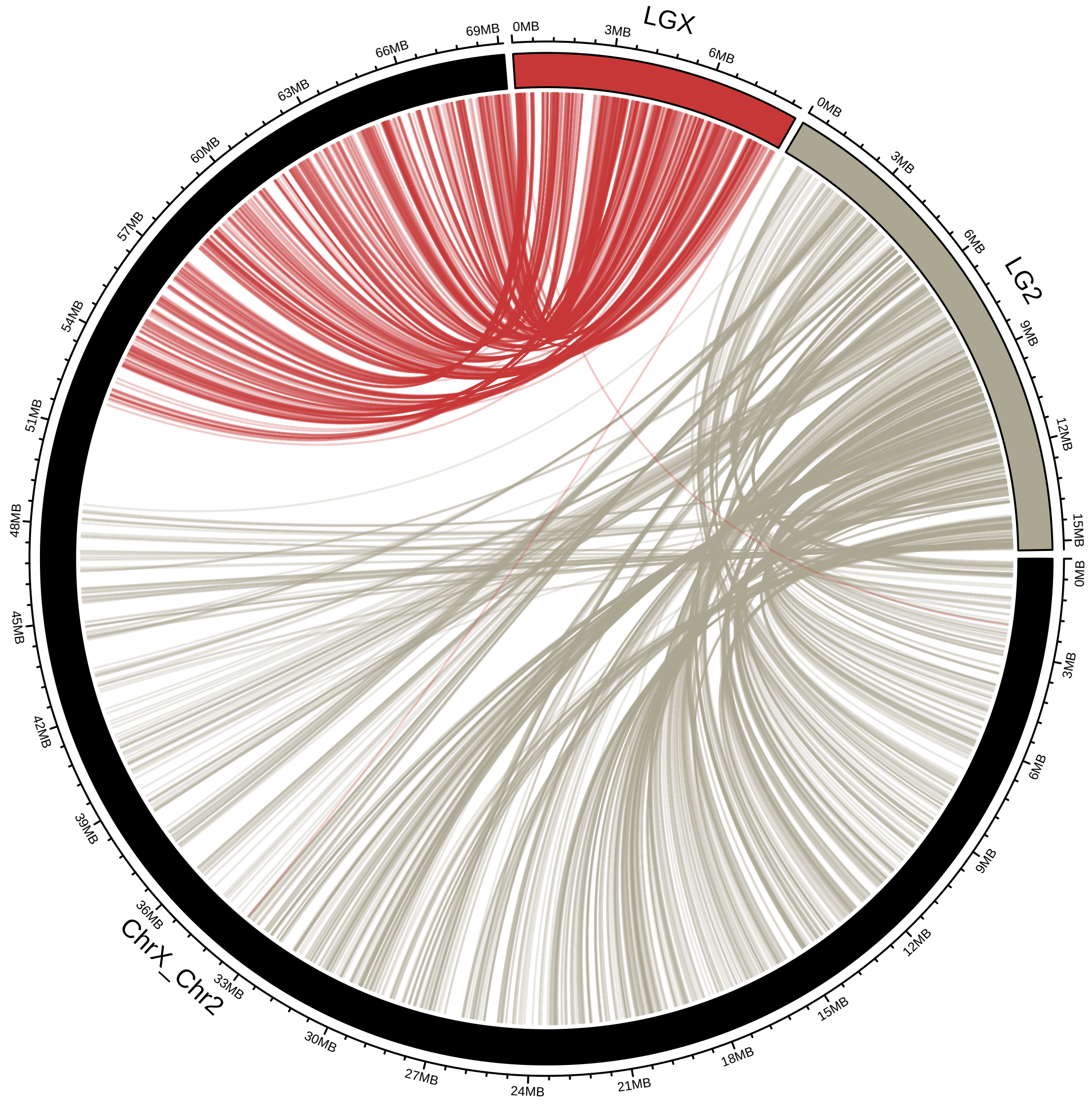

### Supplementary Figure 3

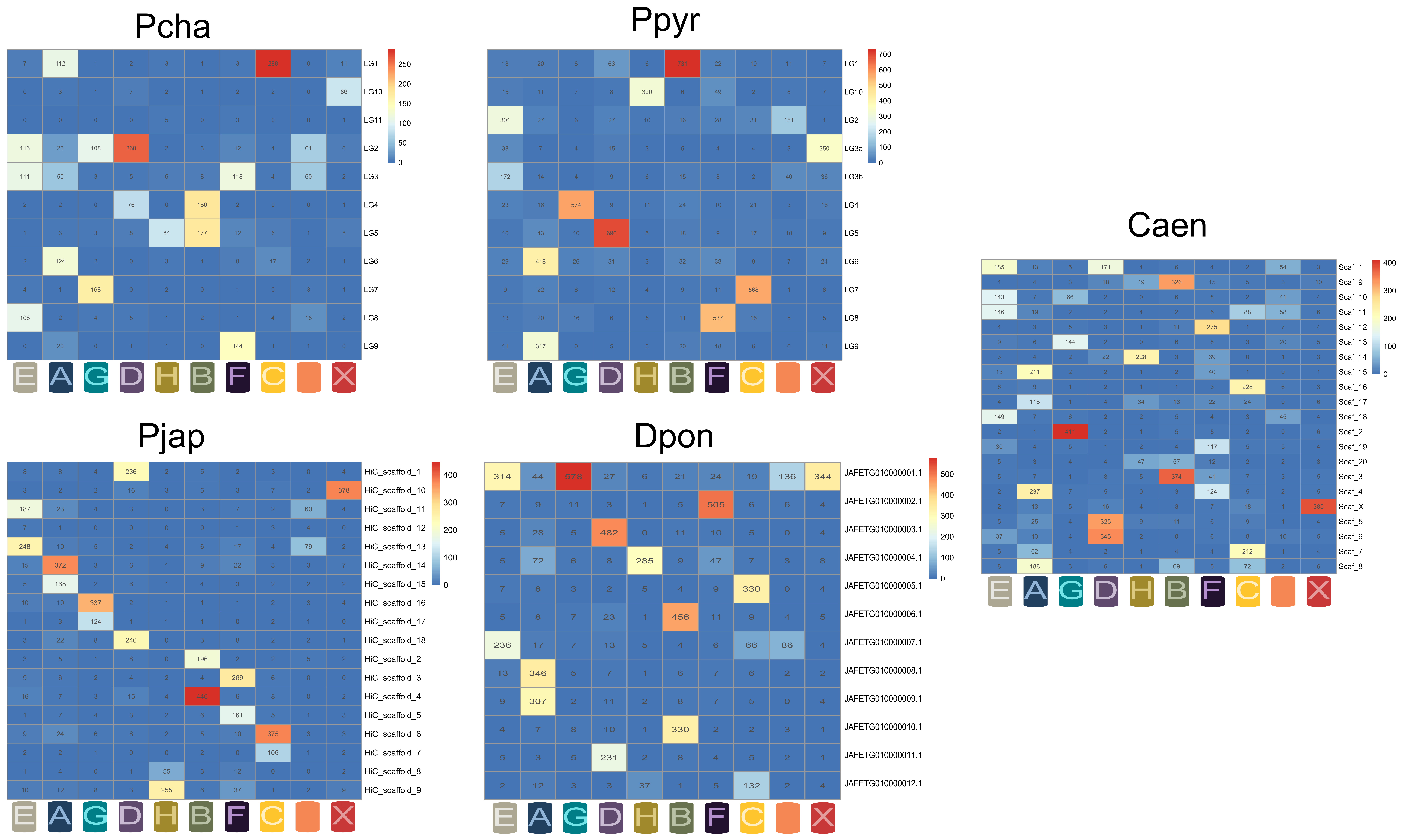

### Supplementary Figure 4

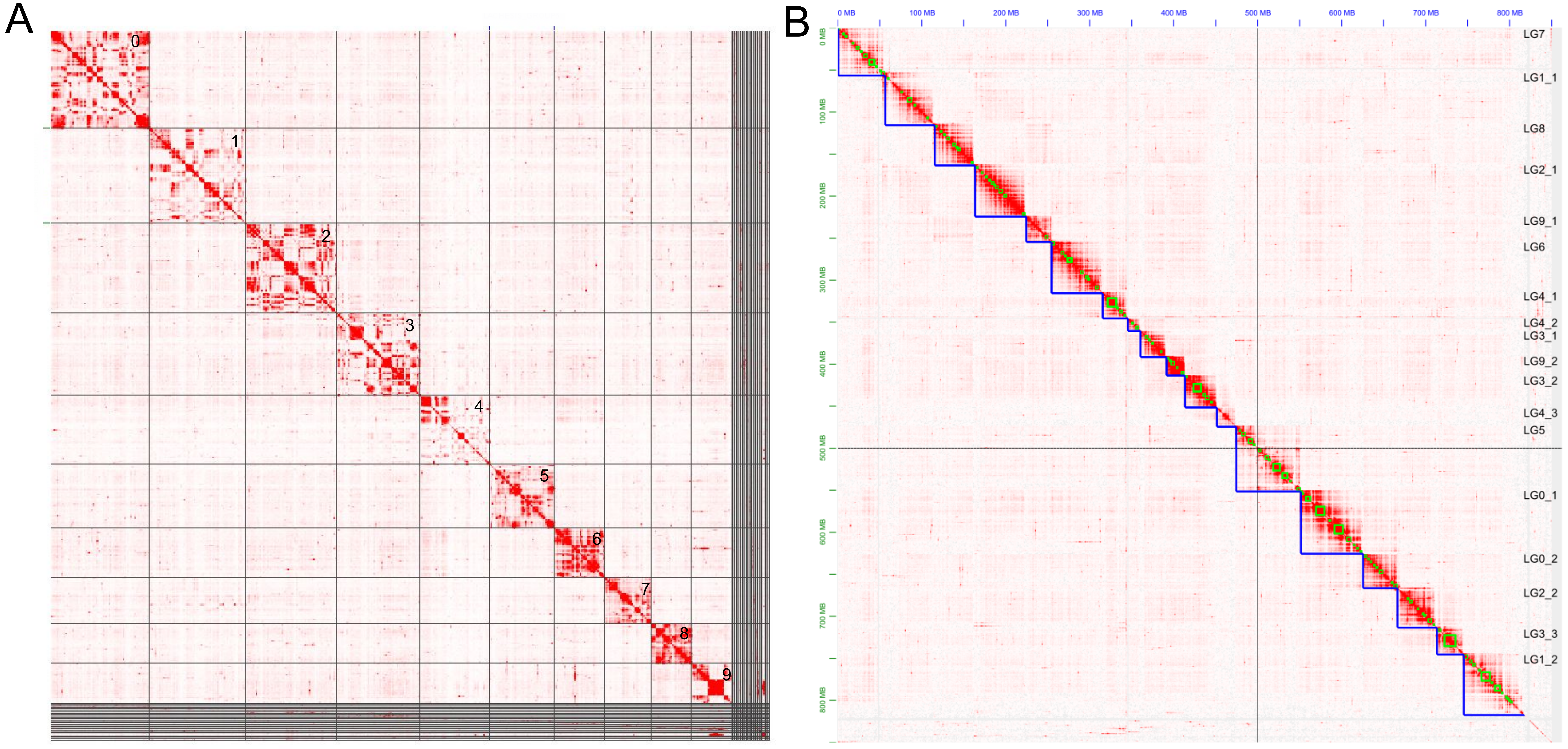
